# Evolution of beta-lactam resistance causes fitness reductions and several cases of collateral sensitivities in the human pathogen *Haemophilus influenzae*

**DOI:** 10.1101/2025.05.02.651845

**Authors:** Sabine Petersen, Margo Diricks, Christian Utpatel, Hinrich Schulenburg, Matthias Merker

**Affiliations:** Evolution of the Resistome, Research Center Borstel, Borstel, Germany; Molecular and experimental Mycobacteriology, Research Center Borstel, Borstel, Germany; German Center for Infection Research, Partner site Hamburg-Lübeck-Borstel-Riems, Borstel-Lübeck, Germany; Evolutionary Ecology and Genetics, Kiel University, Kiel, Germany; Antibiotic Resistance Group, Max-Planck Institute for Evolutionary Biology, Ploen, Germany

**Keywords:** Haemophilus influenzae, ampicillin resistance, cephalosporine resistance, evolution of antimicrobial resistance, collateral effects

## Abstract

The evolution of antimicrobial resistant (AMR) pathogens poses a global health threat and it is unclear if evolutionary trajectories to resistance lead to predictable phenotypes. We analyzed AMR-evolution in the human pathogen *Haemophilus influenzae* in controlled evolution experiments with increasing concentrations of either ampicillin, cefotaxime, or ceftriaxone. We isolated 315 clones from different time points of six independent experiments and characterized changes in genome sequences, bacterial fitness, and minimum inhibitory concentrations (MICs) to 14 antibiotics. Resistance evolution under ampicillin and cefotaxime was mainly driven by mutations in the *ftsI* gene, encoding the penicillin-binding protein 3. However, ceftriaxone exposure repeatedly selected for amino acid substitutions in the outer membrane protein P2 (OmpP2). Some OmpP2-mutants reproducibly showed phenotypic heterogeneity for ceftriaxone but also for fluoroquinolones, rifampicin and tetracycline. Bacterial fitness assessments revealed trade-offs between resistance-associated mutations and growth, though no systematic correlation of MIC increase and growth deficit was detected. Over 50% of the selected clones became more susceptible to aminoglycosides, clarithromycin, and colistin, while some mutation patterns resulted in cross-resistance to meropenem and fluoroquinolones. Overall, beta-lactam antibiotics reproducibly selected mutants with increased MICs, but evolutionary pathways and resulting phenotypes remain unpredictable. These findings highlight the complexity of resistance evolution and suggest that future AMR treatment strategies may need to consider strain-specific and collateral effects more closely.

## Introduction

Antimicrobial resistance (AMR) is a major public health challenge globally (1). The WHO has identified several priority pathogens to inform research and development as well as public health interventions to combat the AMR crisis (2). One of these priority pathogens are ampicillin resistant *Haemophilus influenzae* strains. *H. influenzae* is a common Gram-negative opportunistic pathogen residing in the human upper respiratory tract (3,4). The bacterium can cause a range of infections, from mild illnesses like sinusitis and otitis media to severe invasive diseases such as meningitis and bacteremia, particularly in vulnerable populations (5). Despite the successful implementation of a vaccine against *H. influenzae* serotype b strains, non-typeable strains (NTHi), i.e., strains lacking a polysaccharide capsule, have become prevalent (6) and evolved resistance to aminopenicillins such as ampicillin and amoxicillin in many world regions (7–9).

While amoxicillin-clavulanate and ampicillin-sulbactam are used worldwide to overcome beta ⍰ lactamase-mediated resistance (predominantly TEM-1 and ROB-1), third-generation cephalosporins such as cefotaxime and ceftriaxone also effectively bypass these enzymes (10,11) and are increasingly favored in clinical practice (12). This is due to their favorable tolerability and their ability to achieve high serum and cerebrospinal fluid concentrations, making them especially suitable for the treatment of severe infections like sepsis and meningitis caused by *H. influenzae* (13). However, resistance to beta-lactams can also evolve in beta-lactamase negative strains via alterations in the penicillin binding protein 3 (PBP3), encoded by the *ftsI* gene (14). Mutations in *ftsI*, particularly in the transpeptidase domain of PBP3, reduce the binding affinity for different beta-lactam antibiotics (15). Current understanding is limited regarding whether resistance evolution in *H. influenzae* follows identical genetic and phenotypic trajectories across different beta-lactam classes, and whether such evolutionary pathways are reproducible. In particular, it is unclear if evolution consistently leads to cross-resistances against multiple beta-lactam antibiotics, or if variability in resistance mechanisms leads to diverse outcomes. Here, laboratory-based evolution experiments provide a powerful approach to address these questions (16). By simulating selective pressures *in vitro*, these experiments allow for tracking genetic changes and phenotypic adaptations as resistance evolves. In addition, this experimental approach facilitates an assessment of evolved collateral effects, such as cross-resistance or collateral sensitivity (i.e., variants with evolved resistance against certain antibiotics show an increase in susceptibility to other antibiotics), which may inform about epistatic interactions, and possibly new treatment regimens against resistant strains (17).

Therefore, the aim of the current study was to enhance our understanding of the predictability of resistance evolution and the resulting collateral effects. In particular, we investigated the evolutionary pathways leading to resistance against either ampicillin, cefotaxime, or ceftriaxone in *H. influenzae* using a multi-step *in vitro* evolution approach. By subjecting the reference strain *H. influenzae* Rd KW20 to increasing concentrations of these antibiotics, we tested whether evolution results in identical or divergent genetic trajectories and phenotypes. We employ short- and long-read sequencing technologies to map mutations across the genome and assess their impact on minimum inhibitory concentrations (MICs) for different antibiotics. Furthermore, we evaluated whether the evolution of resistance is reproducible by examining the consistency of mutation patterns and collateral effects on other antibiotics.

## Methods

### Bacteria and culture methods

The beta⍰ lactamase negative reference strain *H. influenzae* Rd KW20 and evolved mutant strains were routinely grown from ⍰80°C freezer stocks on chocolate agar (Mueller-Hinton broth with 5 g/L yeast extract, 17 g/L agar, 5% defibrinated horse blood and 20 mg/L beta-nicotinamide adenine dinucleotide (beta-NAD)) at 37°C in a humid atmosphere containing 5% CO_2_. Colonies grown on chocolate agar served as inoculum for liquid cultures in Mueller-Hinton broth supplemented with 5 g/L yeast extract, 20 mg/L beta-NAD and 15 mg/L hemin, hereafter referred to as *Haemophilus* test medium (HTM), grown at 37°C and shaking at 100 or 200 rpm. Liquid cultures were routinely plated on HTM agar (formulation of HTM with 17 g/L agar added).

### Antibiotic susceptibility testing

To determine the antibiotic starting concentrations for the multi-step evolution experiment, MICs of ampicillin, cefotaxime and ceftriaxone were tested for the reference strain using broth microdilution methodology according to EUCAST protocol, but using HTM. Ampicillin sodium salt, cefotaxime sodium salt, and ceftriaxone disodium salt hemi(heptahydrate) were purchased from Sigma-Aldrich.

Gradient diffusion strips (Liofilchem® MIC Test Strips) on Mueller Hinton fastidious agar (MH-F, BD) were routinely used for all isolated single clones evolved in the evolution experiment to determine MICs of ampicillin, cefotaxime and ceftriaxone. To check for collateral effects, 17 clones (3–4 per evolved population, selected based on differing mutational patterns and/or distinct beta-lactam MICs) were additionally tested for clarithromycin, kanamycin, colistin, vancomycin, levofloxacin, rifampicin, amikacin, tetracycline, gentamicin and meropenem susceptibility using Liofilchem® MIC Test Strips. Susceptibility interpretation was performed according to EUCAST breakpoints v. 13.0 2023, which are 1 mg/L for ampicillin, 0.125 mg/L for cefotaxime, and 0.125 mg/L for ceftriaxone.

### Multi-step evolution experiment

Populations of the susceptible reference strain *H. influenzae* Rd KW20 were evolved under antibiotic pressure (ampicillin, cefotaxime, ceftriaxone) in HTM, starting with an antibiotic concentration of 0.5xMIC of the reference strain (determined using broth microdilution methodology). One population was evolved in the absence of any antibiotic pressure (control). Evolution in the presence of ceftriaxone was performed three times to investigate reproducibility of resistance evolution.

A volume of 20 µL of single colony resuspension (one single colony of *H. influenzae* Rd KW20 grown on chocolate agar resuspended in 100 µL of HTM) was used to inoculate 2-mL cultures containing either 0.125 mg/L of ampicillin, 0.008 mg/L of cefotaxime, 0.002 mg/L of ceftriaxone (0.002 mg/L) or no antibiotic (1^st^ passage) (Supplementary Figure S1).

The evolving 2-mL cultures were grown in 12-well microtiter plates at 37°C, 100 rpm and 5% CO_2_ for four days per cycle, with daily 100-fold dilution into 2 mL of fresh HTM. At each passage, one culture was grown at the same antibiotic concentration as in the previous passage (back-up culture), while three additional cultures were inoculated containing twice the antibiotic concentration of the previous passage. Whenever at least one of the cultures with higher antibiotic concentration showed visible growth after 24 h, this culture was used to inoculate the next passage. This approach allowed for a stepwise increase in antibiotic concentration throughout the experiment without extinction. The highest antibiotic concentration that supported visible growth after 20 passages was recorded as the final concentration: 1 mg/L for ampicillin, 0.125 mg/L for cefotaxime, and 0.064 mg/L for ceftriaxone. At the end of each four-day cycle, 20 µL of culture were spotted onto chocolate agar and incubated for three days at 37°C and 5% CO_2_. After incubation, the biomass of the spotted cultures was resuspended in 1 mL of HTM, and 20 µL of this resuspension were used to initiate the next cycle. This process was repeated for a total of five cycles adding up to twenty passages in liquid medium. In the final cycle, cultures were not spotted onto chocolate agar but directly processed. Glycerol stocks (HTM with 20% glycerol) of each passage were stored at -80°C.

Single clones were isolated from each passage by dilution streaking of the respective glycerol stocks on chocolate agar. In general, two clones were isolated per passage. However, whenever the antibiotic concentration was increased compared to the previous passage, five clones were isolated instead. Isolated clones were grown in 3 mL of HTM to determine MICs of ampicillin, cefotaxime, and ceftriaxone using gradient diffusion strips (Liofilchem® MIC Test Strips) and to isolate DNA using the DNeasy UltraClean Microbial Kit (Qiagen) for DNA sequencing.

### Whole genome sequencing

Short-read DNA-libraries were prepared from extracted genomic DNA with a modified Illumina Nextera XT library kit protocol as described previously (18). Libraries were sequenced with 150 bp paired-end reads on an Illumina NextSeq 2000 (Illumina, San Diego, CA, USA). Short-read data (fastq files) are available in the European Nucleotide Archieve (ENA) under the BioProject PRJEB86018. Accession numbers are available in Supplementary Table S1. Mutations, i.e., single nucleotide polymorphisms (SNPs), short insertions and deletions, were called with a reference mapping pipeline (19). Briefly, short reads were mapped to the reference genome *H. influenzae* Rd KW20 (ATCC 51907) (BioSample SAMN39831887) using the Burrows-Wheeler Aligner (BWA) (20) and allele calling was done with the Genome Analysis Toolkit (GATK) (21). Mutations were called with the following thresholds: 4 reads mapped in both forward and reverse orientation, a minimum of 8 reads with a PHRED score ≥20 and ≥75% allele frequency. Indels in the gene *hdsM* (i.e., repetitive region) were excluded. For subsequent phylogenetic analysis, SNPs were concatenated when at least 95% of all isolates fulfilled the abovementioned thresholds for read coverage, PHRED score and frequency at individual genome positions.

Long-read sequencing was performed with the PacBio Sequel II system (Pacific Biosciences, CA, USA) for selected clones at different timepoints from each evolution experiment to identify structural genomic changes. DNA libraries were prepared with the SMRTbell Express Template Prep Kit 2.0 with barcoded adapters from IDT (Integrated DNA Technologies, USA). Alignment files (.bam) are available in the ENA under the BioProject PRJEB86018. De-novo genome assembly was performed using the PacBio SMRTlink software v9.0 and its “Microbial Assembly” application, with the genome length set to 1.8 Mb and a seed coverage of 30. For the resulting assemblies, the same origin was set using the first 50 bp of dnaA. This sequence was identified using the Seqkit toolkit (Seqkit v2.9.0), and the origin was adjusted accordingly. Subsequently, each assembly was aligned to the reference genome using progressiveMauve (Mauve v2.4.0), and the detected variations were extracted from the alignment. Indels of at least four bases in length were further examined through visual inspection of the assembly and reference mapping in these regions to characterize the mutation more precisely (tandem repeat variation, inversion, deletion, insertion).

Long-read sequencing detected deletions and inversions in *ompP2*, which were not detected using Illumina sequencing. To check for presence of structural changes affecting *ompP2* the BWA alignment files of all 315 clones were checked manually for variations in *ompP2* using Geneious Prime v2024.0.5 (https://www.geneious.com/).

### Phylogenetic and Bayesian analysis

Fasttree v.2.1.9 (22) was used to calculate phylogenetic trees for each evolution experiment with a general time reversible (GTR) substitution model and Gamma20 likelihood optimization to account for evolutionary rate heterogeneity among sites and 500 bootstrap replicates. Figtree v.1.4.4 (https://github.com/rambaut/figtree/releases) was used to root phylogenetic trees at midpoint and iTol v.6.8.1 (https://itol.embl.de/) was used for visualization. A phylogenetic molecular clock signal was first assessed with TempEst v1.5 (23) by the correlation of sampling day and root-to-tip distance. We further performed a permutation-based linear regression analysis using the lmPerm v2.1.0 R package, to tests the significance of the regression model without assuming normality by computing values through randomized permutations of the data. Mutation rates were calculated with BEAST2.5 (24) for all evolution experiments, except one replicate experiment exposed to ceftriaxone with a poor correlation of SNPs and sampling time, and a non-converging Markov-Chain-Monte-Carlo sampling. We employed a coalescent exponential population model, log normal clock rate, a strict molecular clock, tip dating (day of the experiment), a GTR substitution model, 20 million iterations with 10% burn-in, and sampling every 2,000 traces/trees. Inspection of BEAST log files with Tracer v1.7.2 showed an adequate mixing of the Markov chains and all parameters were observed with an effective sample size (ESS) > 200.

### Competitive growth assay

Pairwise competitive growth assays were performed to estimate the *in vitro* relative fitness of mutants evolved in the multi-step evolution experiment compared to the parental strain (*H. influenzae* Rd KW20). Investigated clones were selected to represent a diverse range of mutational patterns. Briefly, five single colonies grown on chocolate agar served as inoculum for 3 mL precultures that were grown until mid-exponential phase (Supplementary Figure S2). Pre-cultures were diluted to an OD_600_ of around 0.025. Equal volumes (1.5 mL) of diluted mutant and parental strain cultures were added in 15 mL falcon tubes. Co-cultures were grown in triplicate for 4 h at 37°C and shaking at 200 rpm. Initial and final colony forming units (CFU) of the mutant and parental strain were determined by standard plate counting on HTM agar (total CFU) and on HTM agar containing 0.016 g/mL cefotaxime (mutant CFU), respectively. Even though co-cultures were inoculated to a 1:1 ratio of the OD_600_ of the two competing strains, initial CFU counts sometimes differed between both strains. Initial CFU ratios of the parental strain to that of the mutant strain of 1:0.5 to 1:2 were accepted, as the formula used below accounts for unequal starting CFUs. Relative fitness was assessed as the ratio of the Malthusian parameter of the mutant to that of the parental (reference) strain. The following formula considering population sizes of the mutant and the parental strain was

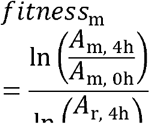

used for the calculation of fitness of each strain in relation to the parental strain (25).

The formula calculates the fitness (*fitness*_m_) of the mutant strain relative to the reference strain (Rd KW20). Here, *A* represents the population size, with subscripts m for the mutant strain and r for the reference strain. The time points 0h and 4h indicate the population size at inoculation and after 4 hours, respectively. A fitness of 1 indicates no fitness cost, whereas a ratio greater than or less than 1 indicates increased or decreased fitness compared to the parental strain, respectively.

### Crystal structures

The crystal structure of the transpeptidase domain (amino acid 254-610) of PBP3 of *H. influenzae* was obtained from the RCSB Protein Data Bank (entry ID 6HZO deposited by Bellini et al.). Detected PBP3 substitutions were visualized in this crystal structure using PyMOL v3.0 (https://www.pymol.org/). Ampicillin and cefotaxime were superimposed using crystal structures of PBP3 of *Mycobacterium tuberculosis* (entry ID 6KGW) and *Staphylococcus aureus* (entry ID 3VSL). Structure of full PBP3 (entry ID AF-P45059-F1-v4) was obtained from AlphaFold Protein Structure Database.

AlphaFold Server powered by AlphaFold 3 (26) was used to predict structures of wild-type and mutated Outer membrane protein P2 (OmpP2). Predicted structures were visualized using PyMOL v3.0.

### Statistical analyses

The association between specific mutations and a reduction in susceptibility was analyzed using Mann-Whitney U tests. For this purpose, the ampicillin, cefotaxime, and ceftriaxone MIC values of two sets of evolved clones were compared. The clones in the two sets differed genetically only by the mutation or mutation combination under investigation (based on short-read sequencing approach and the manual check of *ompP2* mutations in the BWA alignments), aside from detected tandem repeat variations in the *hsdM* gene. For each clone set, a minimum of three genetically identical clones was required. To minimize the risk of false-positive results due to multiple testing, *p*-values were adjusted using the Benjamini-Hochberg correction. Given that all clones were derived from the same ancestral strain, we acknowledge the possibility of non-independence among observations and interpreted statistical outcomes with caution.

A one-sample t-test (considering the Malthusian parameters ln(*A*_m, 4h_/*A*_m, 0h_) and ln(*A*_r, 4h_/*A*_r, 0h_)) with Benjamini-Hochberg correction for multiple testing was used to assess the significance of fitness gain/loss of mutants compared to the reference strain. Pairwise t-tests with Benjamini-Hochberg correction were conducted to identify significant differences between fitness values of clones isolated from the same evolving population.

A permutation test was conducted to evaluate the association between average fitness and antimicrobial susceptibility (MIC) to ampicillin, cefotaxime, and ceftriaxone. For each antibiotic, the observed correlation between relative fitness value and MIC was calculated. The null distribution was generated by randomly shuffling the MIC values within each group and recalculating the correlation for 10,000 permutations. The *p*-value was determined by comparing the observed correlation to the null distribution, assessing the statistical significance of the observed association.

## Results

### Evolution of beta-lactam resistance is mainly driven by *ftsI* and *ompP2* mutations

We exposed the reference strain *H. influenzae* Rd KW20 in a multi-step evolution experiment to increasing concentrations of the three beta-lactam antibiotics ampicillin, cefotaxime, and ceftriaxone. Bacterial populations evolved over 20 transfers in individual experiments, which is equivalent to approximately 200 generations. Overall, we investigated the genomes and MICs of 315 clones, isolated across time from six independent evolution experiments including a drug-free control and three replicate experiments with ceftriaxone. We identified between 11 and 25 mutations (including SNPs and indels) in individual experiments over 20 days of exponential growth, resulting in a linear increase in the number of mutations across time. SNP-based phylogenies and root-to-tip linear regressions (R2 0.31-0.86) indicated moderate to strong support of a constant molecular clock in all evolution experiments, except for one replicate exposed to ceftriaxone (R2 = 0.11) (Supplementary Table S3). Based on a Bayesian coalescent model, assuming a strict molecular clock and exponential population growth, the mutation rates ranged from 1.0 × 10^−7^ to 2.1 × 10^−7^ substitutions per site per day.

The MICs against the three beta-lactam antibiotics of individual clones did not change significantly in the drug-free control experiment (Supplementary Figure S3). In the drug-free control, measured MIC differences were within the range of technical variation, i.e., one doubling dilution compared to the MICs of the parental strain Rd KW20, which were 0.19 mg/L for ampicillin, 0.008 mg/L for cefotaxime, and 0.003 mg/L for ceftriaxone. We also did not detect any mutations in the main drug target *ftsI* in the evolved control population (Supplementary Table S1).

In contrast, exposure to ampicillin, cefotaxime and ceftriaxone resulted in a pronounced increase of beta-lactam MICs (Figure 1). Specifically, exposure to ceftriaxone led to strong MIC increases, including in one clone an MIC increase of up to 64-fold against ceftriaxone relative to the ancestor strain. Exposure to ampicillin resulted in more moderate MIC increases against all three beta-lactam antibiotics. Furthermore, populations exposed to third-generation cephalosporins showed a more pronounced MIC increase against ceftriaxone and cefotaxime as compared to ampicillin.

**Figure 1.**
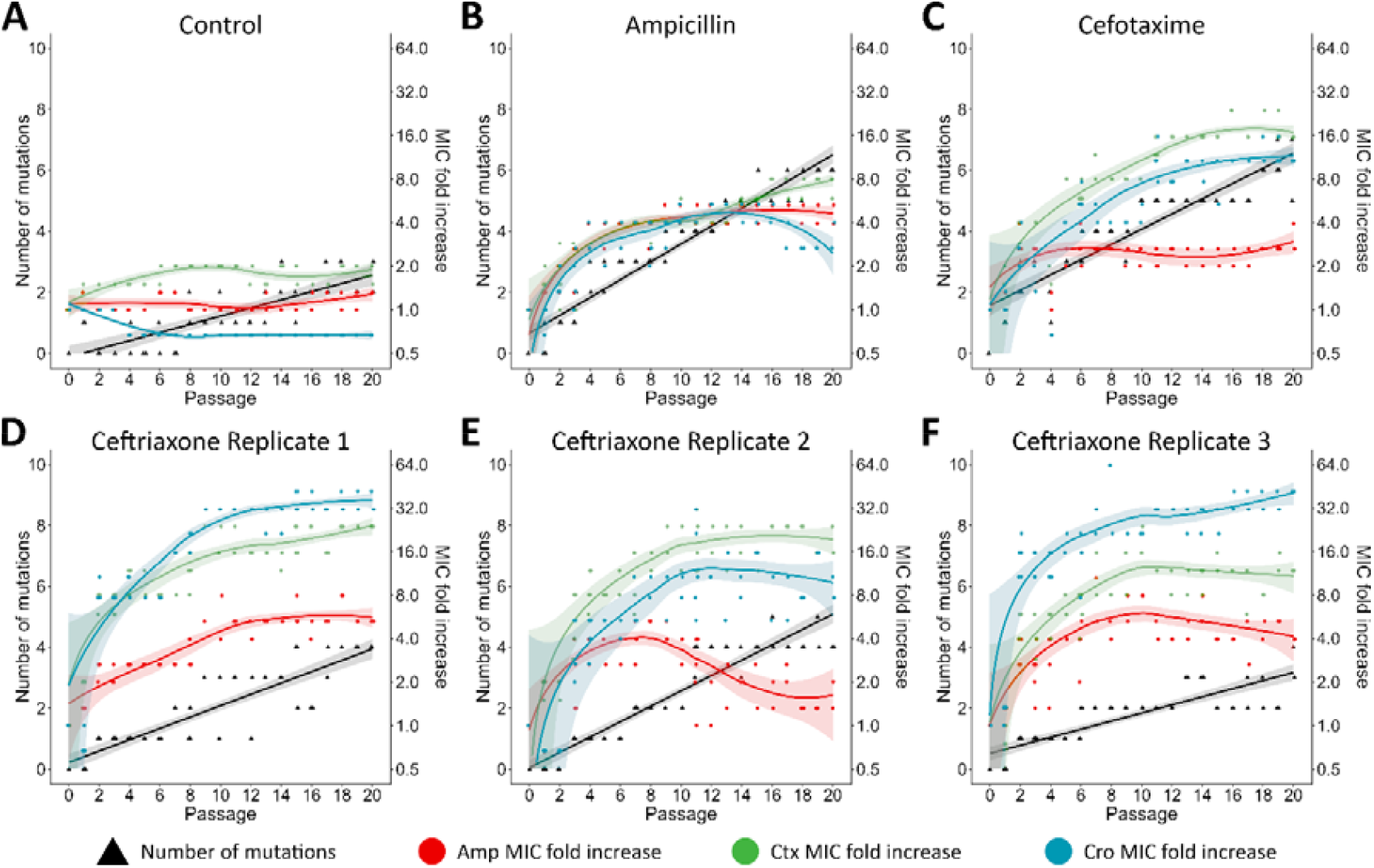
Increasing number of mutations and changes of beta-lactam MICs in six evolution experiments. Number of mutations (black) and minimum inhibitory concentration fold-increase compared to the parental strain Rd KW20 for ampicillin (Amp, red), cefotaxime (Ctx, green), and ceftriaxone (Cro, blue) in selected clones over 20 passages (two or five clones per passage). Bacterial population A - without antibiotic exposure and exposed to increasing concentrations of B - ampicillin, C - cefotaxime, D - ceftriaxone replicate 1, E - ceftriaxone replicate 2, F - ceftriaxone replicate 3. Shaded areas represent the 95% confidence interval.

Clinical resistance against cefotaxime (EUCAST breakpoint: > 0.125 mg/L) was observed for 4/55 (7.3%) clones evolved under cefotaxime exposure and for 28/169 (16.6%) clones evolved in three evolution experiments under ceftriaxone exposure (Figure 2, Supplementary Figure S3). Exposure to ceftriaxone further selected 1/169, (0.6%) ceftriaxone resistant (EUCAST breakpoint: > 0.125 mg/L) clone and 10/169 (5.9%) ampicillin resistant (EUCAST breakpoint: > 1 mg/L) clones. Cross-resistance against ampicillin and cefotaxime was observed for five clones evolved under ceftriaxone exposure (Supplementary Figure S3). Overall, 20 days exposure to ampicillin did not select for clinical resistant clones against any of the three tested drugs.

**Figure 2.** Maximum likelihood phylogeny of *H. influenzae* Rd KW20 clones evolved in the presence of ampicillin (A. S3 clones), cefotaxime (B. 56 clones), and ceftriaxone (C, 57 tones). Amino acid substitutions in the penicillin binding protein 3 (PBP3) and presence of mutations in specified genes as well as minimum inhibitory concentration (MIC) changes over time are color coded. Strains that exceeded the clinical breakpoint for ampicillin/cefotaxime/ceftriaxone according to EUCAST breakpoints are marked with a white star at the MIC heatmap.

As expected, increasing MICs against the beta-lactam antibiotics were associated with mutations in *ftsI* (Figure 2, Supplementary Figure S3). Out of 136 evolved clones that exhibited at least one *ftsI* mutation, 18 were resistant according to EUCAST breakpoints for cefotaxime, but none were resistant to ampicillin or ceftriaxone. Specifically, we identified combinations of the following five amino acid (amino acid) substitutions in individual clones in the transpeptidase domain: p.Thr332Ile (c.968C>T), p.Thr443Ala (c.1327A>G), p.Ala530Ser (c.1591G>T), p.Asp551Tyr (c.1652G>T), and p.Ala561Glu (c.1684C>A). Exposure to both ampicillin and cefotaxime, selected for p.Ala530Ser. Under cefotaxime conditions p.Ala530Ser, p.Thr443Ala, and p.Thr332Ile evolved in a stepwise manner. The substitutions p.Asp551Tyr and p.Ala561Glu were detected independently in separate clones, each arising contemporaneously (detected in 3^rd^ passage) in one population exposed to ceftriaxone. The detected *ftsI* mutations were significantly associated with MIC increase of beta-lactams (Supplementary Table S4). In a crystal structure model of PBP3, all five *ftsI* mutations were either in proximity to other well-known resistance-associated amino acid positions (Ile449, Arg517, Asn526), or close to the binding site of the antibiotics (Figure 3).

**Figure 3.**
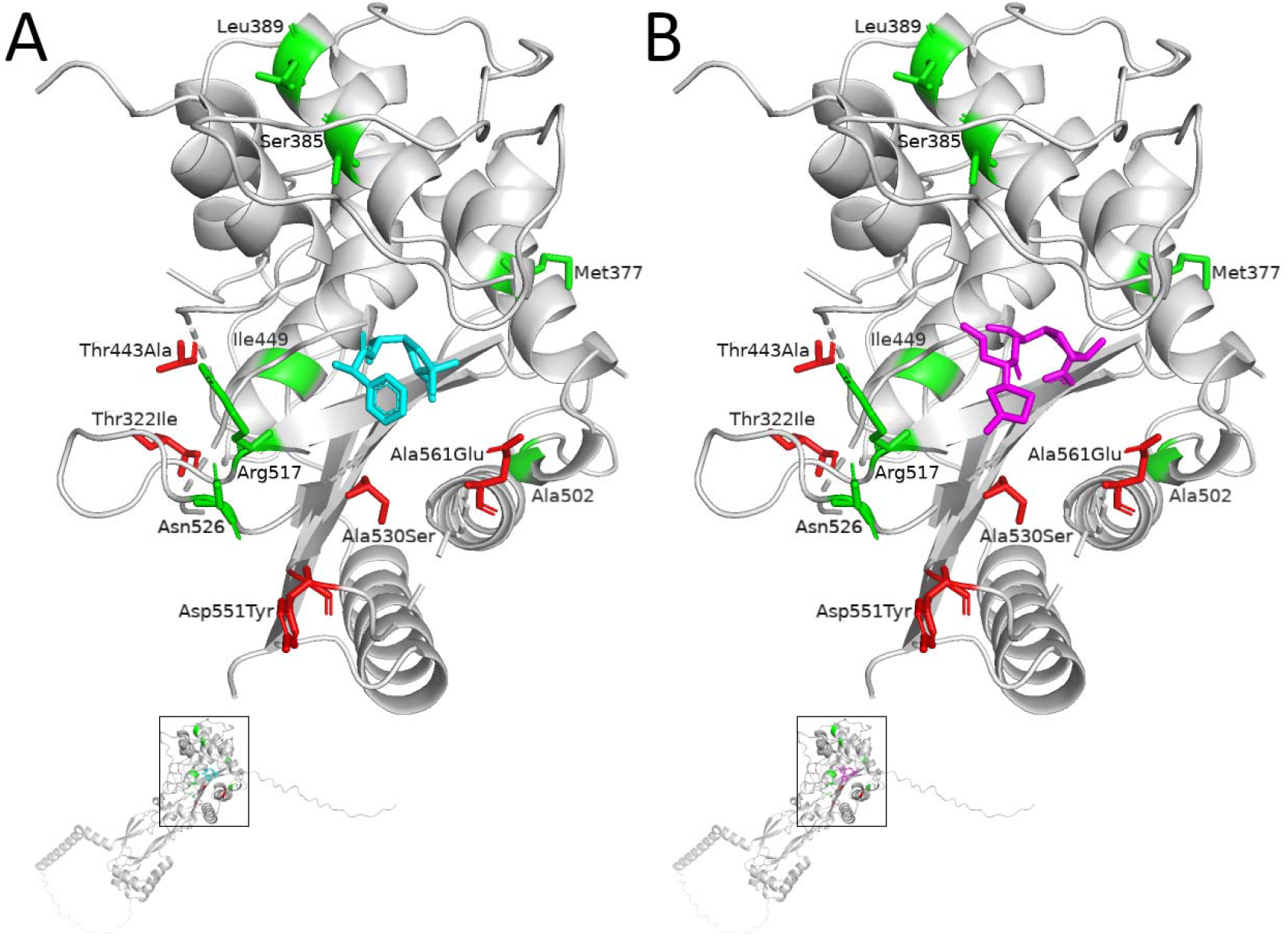
Crystal structure of transpeptidase domain (residues 254-610) of the penicillin binding protein 3 (PBP3) of *H. influenzae* (Bellini et al. [2019]) with superimposed ampicillin (A, colored in cyan) and cefotaxime (B, colored in magenta) molecules. *In vitro* selected amino acid substitutions in PBP3 Thr332Ile, Thr443Ala, Ala530Ser, Asp551Tyr, and Ala561Glu are colored in red. Known positions for amino acid substitutions defining established PBP3 resistance groups are colored in green.

Furthermore, *ftsI* mutations differentially affected the susceptibility against ampicillin, cefotaxime, and ceftriaxone. While we observed a significant MIC increase for ampicillin (2-fold increase, *p*_adj_ = 0.008) and ceftriaxone (3-fold increase, *p*_adj_ = 0.004) associated with amino acid substitution p.Ala530Ser, the secondary amino acid substitution p.Thr443Ala did not significantly affect ampicillin susceptibility but was associated with a two-fold increase of the MIC towards cefotaxime (*p*_adj_ = 0.019) (Supplementary Table S4). Out of the five detected PBP3 substitutions, p.Asp551Tyr showed the highest MIC fold increase towards all three beta-lactams with a maximum 8⍰ fold increase of the MIC of ceftriaxone (*p*_adj_ = 2E-06).

Surprisingly, clones isolated from populations exposed to ceftriaxone (replicate 1 and replicate 3) did not harbor any mutations in *ftsI*. However, we repeatedly selected different mutations in the outer membrane protein P2 (ompP2), i.e., p.Thr126fs (c.376_377insT), p.Val353fs (g.902541_902708del), as well as a structural re-arrangement comprising a genomic inversion starting at codon 353 resulting in p.Gly354fs (g.902541_902708inv). The latter two mutations were not detected with the short-read reference mapping approach but were identified based on the long-read assemblies. However, both mutations could also be detected in clones, for which no long-read sequencing data were available, through a manual check of the BWA alignments. In addition, various tandem repeat variations in different genes could be detected using the long-read sequencing data (Supplementary Table S2). Mutation p.Thr126fs introduces a stop codon at position 129 in *ompP2*, resulting in a truncated protein of only 128 amino acids instead of 359 amino acids (Supplementary Figure S4). The other two mutations starting at codon 353 (i.e., g.902541_902708inv and g.902541_902708del) affect the C-terminus of OmpP2, specifically the last β-strand of the β-barrel (Supplementary Figure S5). The mutation g.902541_902708inv evolved independently in two replicate experiments. All three *ompP2* mutations were significantly (*p*_adj_ < 4E-05) associated with a MIC increase against all three beta-lactam antibiotics with a more pronounced effect for ceftriaxone susceptibility (i.e., 11.5-fold MIC increase, as compared to 2-fold and 3.9-fold increase for ampicillin and cefotaxime, respectively). Overall, 104/315 evolved clones carried ompP2 mutations, of which 10 were resistant to ampicillin, 14 to cefotaxime, and one to ceftriaxone according to EUCAST breakpoints. However, all resistant clones with *ompP2* mutations harbored additional mutations, for example, in the *pnp* gene. Clones with *ompP2* mutations only exhibited highly increased MICs, but these were below the respective EUCAST clinical breakpoints (Supplementary Table S4).

Interestingly, strains harboring the inversion mutation (g.902541_902708inv) resulting in p.Gly354fs were phenotypically heterogeneous concerning beta-lactam susceptibility. Respective strains showed two ellipses for all three beta⍰lactams on the agar plates with gradient diffusion strips (Supplementary Figure S6). The lower MIC corresponded to the MIC observed for the parental strain ± two doubling dilutions.

Besides mutations in *ftsI* and *ompP2*, we detected 13 different mutations in *pnp* (encoding polyribonucleotide nucleotidyltransferase) in clones with and without pre-existing *ftsI* or *ompP2* mutations isolated from populations exposed to cefotaxime, and in two replicated experiments under ceftriaxone. Of those 13 mutations, seven induced a premature stop codon (p.Glu18*, p.Lys47*, p.Glu80*, p.Gln159*, p.Glu220*, p.Glu471*, p.Met601*), three resulted in a frameshift (p.Val423fs, p.Gly444fs, p.Glu580fs) and three were non-synonymous SNPs (p.Ala440Val, p.Gln505Lys, p.Gly626Cys). p.Glu220* and p.Glu580fs were associated with a significant (*p*_adj_ < 0.034) MIC increase for all three beta-lactams in clones with p.Thr126fs in OmpP2 (Supplementary Table S4). The effect of the other detected mutations in *pnp* could not be determined due to an insufficient number of clones carrying the specific mutation pattern. Four different mutations in *rpoB* (encoding DNA-directed RNA polymerase subunit beta), three of them in close proximity (p.Leu149Phe, p.Arg151His, p.Gly154Asp), were detected in clones isolated from populations exposed to ampicillin and ceftriaxone (replicate 2). Furthermore, clones derived from ampicillin, cefotaxime, and ceftriaxone (replicate 2) exposed populations evolved four different mutations in *udk* (encoding uridine kinase) and *spoT* (encoding Guanosine-3’,5’-bis(diphosphate) 3’-pyrophosphohydrolase). Two mutations in *prs* (encoding ribose-phosphate pyrophosphokinase) were detected in ampicillin exposed clones as well as two mutations in Rd_08000 (encoding a putative TonB-dependent receptor) in ceftriaxone exposed clones (replicate 1 and 3). However, only minor effects were observed, with a maximum median MIC fold increase of 3 for mutations in genes other than *ftsI* and *ompP2*.

### Evolved resistant mutants pay a fitness cost under drug-free conditions

To investigate possible fitness costs of specific mutants and identify putative compensatory mutations that minimize the fitness costs imposed by resistance-associated mutations, a competitive growth assay was performed. Most of the evolved clones exhibited a lower fitness compared to the parental strain (Figure 4 and Supplementary Table S1), strongly indicating an evolutionary trade-off between resistance and competitive fitness in the absence of antibiotics. However, three strains (E_0.5Amp_07a, E_1Amp_17a and E_0.032Ctx_07a) did not show a fitness/growth defect. One of these clones (E_0.5Amp_07a) harbored putative beta⍰lactam susceptibility decreasing mutations in *ftsI* and *rpoB* (27–29) accompanied by a mutation in *udk*. Another clone (E_1Amp_17a) showed the same mutation pattern but with additional mutations in *rpoA, plsB*, and *prs*.

**Figure 4.**
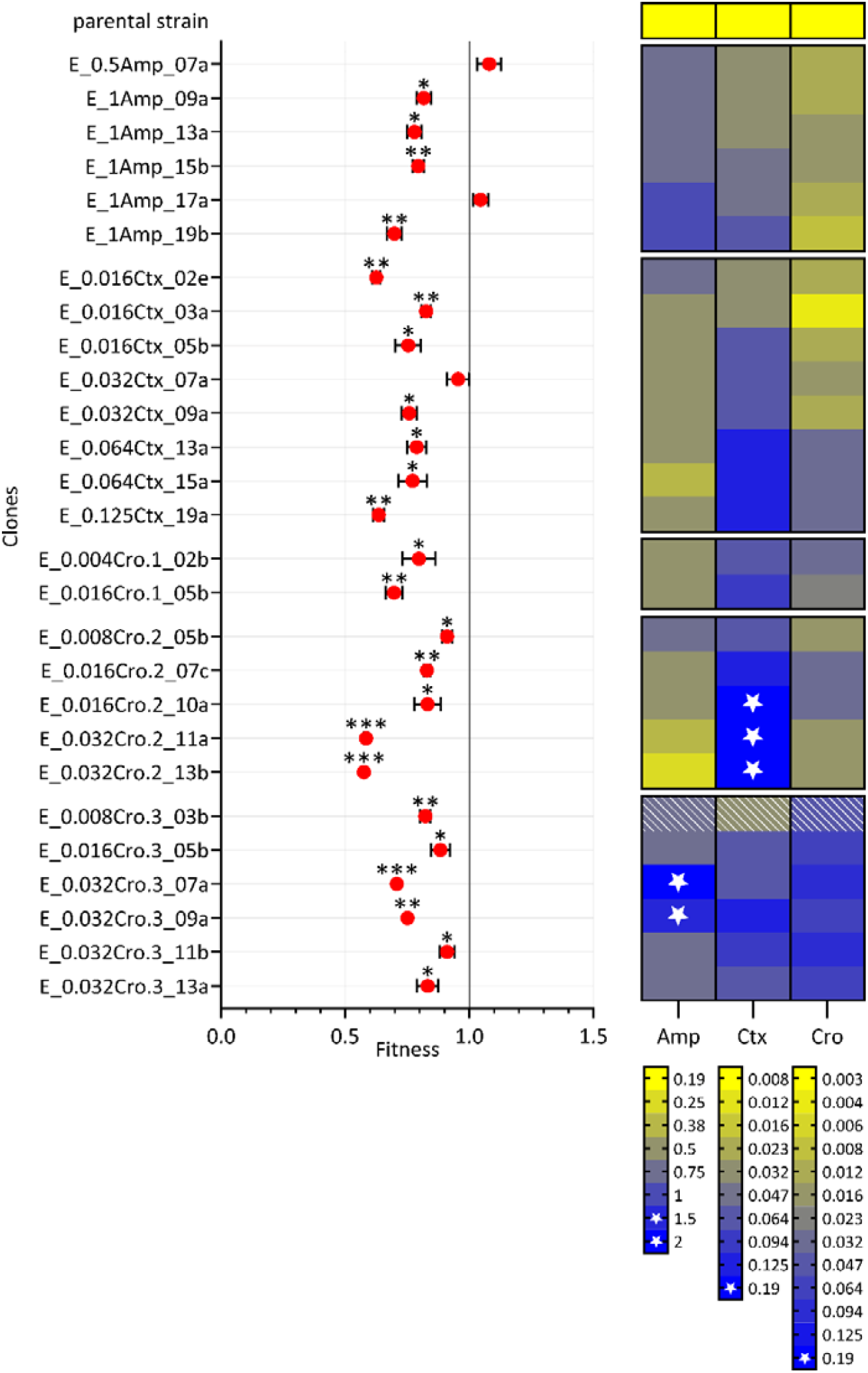
Bacterial fitness of selected clones derived from evolving populations exposed to ampicillin, cefotaxime, and ceftriaxone. Bacterial fitness measured as relative growth difference between the ancestral strain (*H. influenzae* Rd KW20) and the selected clone. A relative fitness of 1 indicates no fitness cost, whereas a value greater or less than 1 indicates increased or decreased fitness, respectively. The black bars represent standard deviation. ^*^ *p*_adj_ < 0.05; ^**^ *p*_adj_ < 0.01; ^***^ *p*_adj_ < 0.001. The heatmap indicates MIC values of ampicillin (Amp), cefotaxime (Ctx) and ceftriaxone (Cro) in mg/L of the respective strain (already presented in Figure 2). Stars indicate clinical resistance above the respective EUCAST breakpoints. Hatched panels indicate phenotypic heterogeneity. Clones that derived from the same evolution experiment are grouped together on the y-axis.

Pairwise t-tests were conducted to compare all clones from a given evolution experiment. Particularly interesting are comparisons between clones that differ by only a single mutation, as these allow for the evaluation of fitness effects of individual mutations within a specific genetic background. Comparison of fitness assay results of E_1Amp_17a and E_1Amp_13a revealed, that the *rpoA* mutation restored wild-type fitness level in the respective mutational background (Supplementary Table S5). Clone E_0.032Ctx_07a exhibits putative beta⍰lactam susceptibility decreasing mutations in *ftsI* and *pnp* (30,31) accompanied by mutated *plsB*. In this case, comparison with fitness assay results of E_0.016Ctx_05b, revealed that evolution of the *plsB* mutation resulted in wild-type fitness in the respective mutational background. Moreover, fitness decrease could directly be addressed to and quantified for specific mutations in *ftsI* (p.Asp551Tyr), *ompP2* (p.Thr126fs, p.Val353fs), and *pnp* (p.Glu580fs, p.Glu220*). Correlating MIC increases and relative fitness of evolved clones revealed a negative correlation between MIC and fitness for cefotaxime (coefficient =-0.45, *p* = 0.019), but no correlation of MICs and fitness for ampicillin and ceftriaxone (Supplementary Table S6).

### Evolved beta-lactam resistance is commonly associated with collateral sensitivity towards aminoglycosides, clarithromycin, vancomycin, and colistin

To assess the expression of collateral effects, we characterized resistance to 14 antibiotics from different classes for 17 clones isolated from the evolution experiments. We identified a consistent pattern of decreased susceptibility for beta-lactam antibiotics (Figure 5). This also includes increased MICs of the carbapenem meropenem. In this case, two clones (E_1Amp_10b and E_1Amp_19b) with increased MICs of ampicillin, cefotaxime, and ceftriaxone expressed cross-resistance to meropenem at above EUCAST breakpoint levels. Surprisingly, two clones evolved under ceftriaxone exposure exhibited increased susceptibility (i.e., collateral sensitivity) to meropenem. These two clones additionally showed collateral sensitivity towards all non-beta-lactam antibiotics, excluding tetracycline. In general, collateral sensitivity is most often expressed towards the three considered aminoglycoside drugs (amikacin, kanamycin, and gentamicin), towards the macrolide clarithromycin, as well as the polymyxin drug colistin, possibly indicating a robust emergence of collateral sensitivity upon beta-lactam resistance evolution in *H. influenzae*.

**Figure 5.**
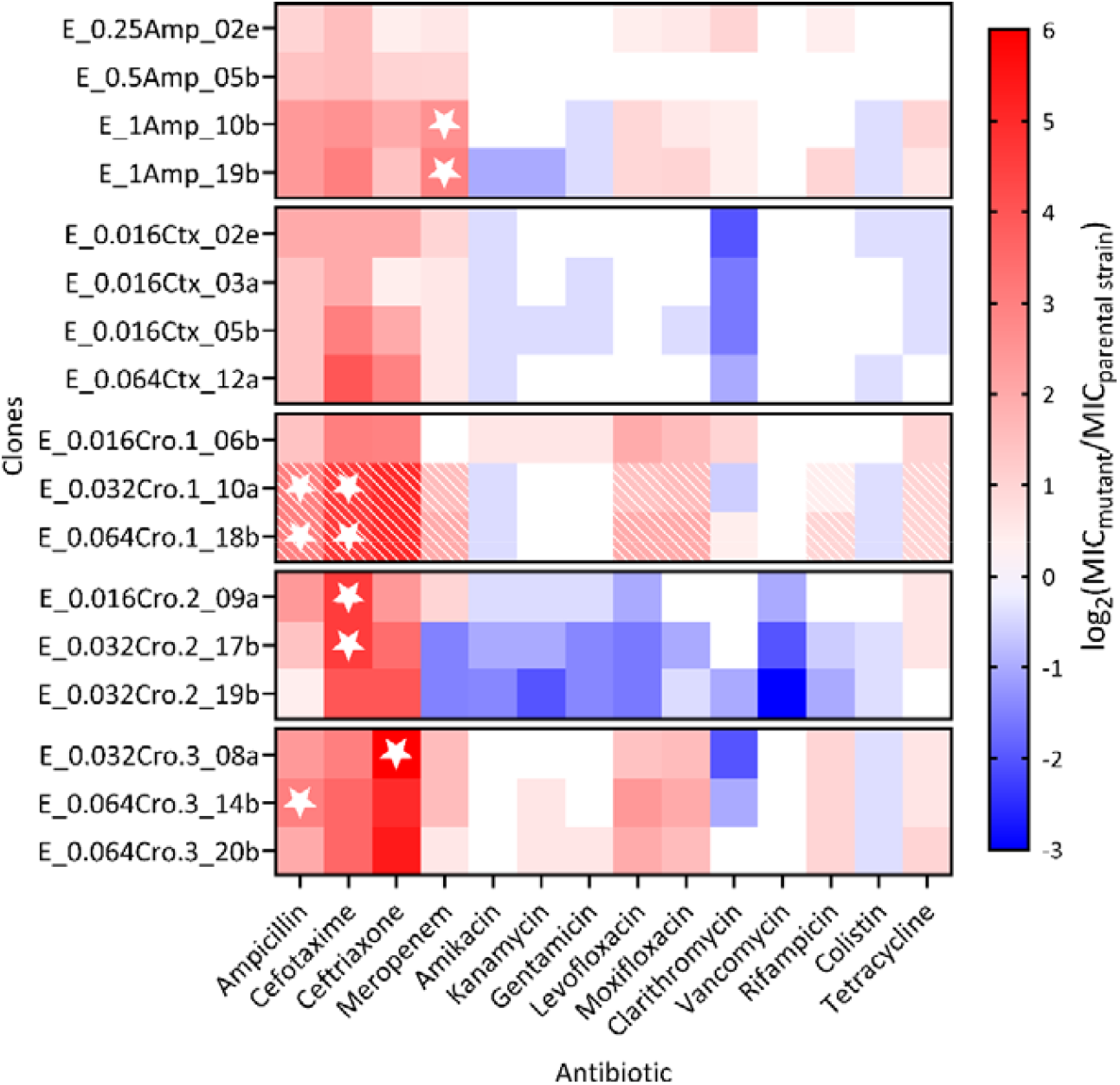
Collateral effects of increased MICs against individual beta-lactam antibiotics. Heat map of log_2_-transformed relative increase in MIC of ampicillin, cefotaxime, ceftriaxone (underlying MIC values already presented in Figure 1), meropenem (beta⍰ lactams), amikacin, kanamycin, gentamicin (aminoglycosides), levofloxacin, moxifloxacin (fluoroquinolones), clarithromycin (macrolide), vancomycin (glycopeptide), rifampicin (ansamycin), colistin (polymyxin) and tetracycline (tetracycline) in mutant clones relative to the MIC of the parental strain *H. influenzae* Rd KW20. Selected clones were evolved in the multi-step evolution experiment. Stars indicate clinical resistance above the respective EUCAST breakpoints (not available for aminoglycosides, clarithromycin, vancomycin, and colistin). Hatched panels indicate phenotypic heterogeneity. Clones that derived from the same evolution experiment are grouped together on the y-axis.

Moreover, the two identified phenotypically heterogeneous OmpP2 mutants consistently expressed phenotypical heterogeneity (i.e., two ellipses with a gradient diffusion test) towards a variety of antibiotics, including next to the beta-lactams also fluoroquinolones, rifampicin and, tetracycline.

## Discussion

This study characterizes the *in vitro* evolution of resistance against three clinically relevant beta-lactam antimicrobials in the opportunistic human pathogen *H. influenzae*. Our results revealed an approximately linear increase in the number of mutations across time, while susceptibility against the antibiotics decreased more steeply during early passages and tended to plateau in the later passages, with dynamics varying across the different beta-lactams and replicates. These increases in resistance are mainly mediated by mutations in previously characterized AMR genes and, in two populations, is additionally associated with the emergence of phenotypic heterogeneity. Generally, evolved resistance against one antibiotic caused significant fitness reductions in drug-free environments and led to altered phenotypic responses to other drug classes, including numerous cases of collateral sensitivity, especially towards aminoglycosides and additional drug classes. Understanding evolutionary pathways and predicting their impact on future treatment regimens will be crucial to reduce bacterial adaptation and selection of antimicrobial resistances.

Respiratory infections with *H. influenzae* are usually treated with an aminopenicillin in combination with a beta⍰lactamase inhibitor (to overcome beta ⍰lactamase-mediated resistance). In case of treatment failure or for severe invasive infections such as sepsis or meningitis, the use of third-generation cephalosporins such as cefotaxime and ceftriaxone is endorsed. However, it is still unclear whether cephalosporin resistance can also develop through alternative evolutionary pathways beyond the well-known PBP3 group III mutations. Furthermore, while resistance to ceftriaxone and cefotaxime is currently rare in clinical isolates (32), understanding potential evolutionary pathways to resistance and collateral effects is crucial for anticipating future threats and optimizing treatment strategies.

To address this, we performed *in vitro* evolution experiments by exposing *H. influenzae* Rd KW20 to increasing concentrations of ampicillin, cefotaxime, and ceftriaxone and monitored genomic changes. Except for p.Thr322Ile, all PBP3 substitutions that were identified in these experiments, were previously reported (in combination with other amino acid substitutions) for patient-derived isolates, and exhibiting decreased susceptibility to cephalosporins and/or ampicillin (p.Thr443Ala (34–37), p.Ala530Ser (34,35,37–40), p.Asp551Tyr (34) and p.Ala561Glu (39)). Notably, none of the PBP3 ampicillin resistance group-defining mutations (p.Met377Ile, p.Ser385Thr, p.Leu389Phe, p.Ile449Val, p.Ala502Val/Thr, p.Arg517His and p.Asn526Lys) evolved in the evolution experiments, although p.Arg517His and p.Asn526Lys are two well-known driver mutations for ampicillin and cefotaxime resistance (41,42). However, PBP3 p.Thr322Ile and p.Thr443Ala, that evolved under cefotaxime exposure, are in close spatial proximity of Asn526 and Arg517, respectively. Interestingly, the resulting structural changes in the beta⍰lactam binding site of PBP3, seem to mainly affect resistance to cephalosporins rather than ampicillin. In *in vitro* evolution experiments with NTHi performed by Jakubu et al.(43), one strain with wild-type *ftsI* evolved p.Thr443Ala in the presence of the cephalosporine cefuroxime. In accordance with our results, the respective population showed only a MIC increase for the cephalosporine, but not ampicillin. Hence, cephalosporins can select for amino acid substitutions that do not affect ampicillin susceptibility.

However, we also observed convergent evolution of p.Ala530Ser under both ampicillin and cefotaxime selection pressure. Respective clones showed significantly increased MICs especially of ampicillin and ceftriaxone demonstrating that amino acid substitution in PBP3 can show collateral effects for beta⍰lactam antibiotics. Further studies, that elucidate to what extent specific *ftsI* mutations and mutation patterns affect third-generation cephalosporin susceptibility as well as information on treatment outcome are essential to evaluate this and improve clinical decision making.

Interestingly, we also observed clones with highly increased beta-lactam MICs that did not harbor any *ftsI* mutations. Ceftriaxone exposure led to evolution of three different *ompP2* mutations resulting in a profound effect on MICs of all three beta-lactam antibiotics. OmpP2 is one of the major outer membrane proteins in *H. influenzae* with a mass of approximately 36 to 42 kDa. The protein functions as a porin with an exclusion limit of about 1.4 kDa (44,45). Studies analyzing the *ompP2* gene of *H. influenzae* reported a high conservation of the 16 β-strands building up the β-barrel, but increased variation and accumulation of SNPs in the eight surface-exposed loop regions (46–49), which can result in antigenic variation (50,51). By expression of OmpP2 and the AcrAB (multidrug efflux pump) encoding genes of *H. influenzae* in an *acrB* and *ompF* knockout strain of *Escherichia coli*, Zwarma et al. provided evidence that sensitivity to beta⍰lactams in *H. influenzae* actually is affected by OmpP2. While the efflux pump AcrAB-TolC actively transports antibiotics from the cytoplasm and periplasm out of the cell, OmpP2 counterbalances this efflux by leaking the drugs back into the cell (52).

Our findings confirm that diverse mutations in *ompP2* can significantly contribute to highly increased beta-lactam MICs in *H. influenzae* (49,53,54). These *ompP2* mutations seem to be particularly selected under ceftriaxone exposure. The *ompP2* frameshift mutation p.Thr126fs detected in this study resulted in a stop codon at amino acid position 129 (instead of position 361) and therefore probably generated a non-functional protein. The other two mutations resulting in p.Val353fs (deletion) and p.Gly354fs (inversion) affected only the last β-strand of the porin, though AlphaFold predicted the mutated sequence to not alter the β-barrel structure. Therefore, decreased influx of beta⍰lactam antibiotics might have resulted from an altered electrostatic profile within the porin channel due to residue properties of the changed AAs (55–57). Most studies on beta⍰lactam resistance in *H. influenzae* focus on PBP3, which makes evaluation of the clinical relevance of *ompP2* mutations and their contribution to treatment failure difficult.

Additional genes, with possibly more moderate effects on ampicillin, cefotaxime, and/or ceftriaxone susceptibility are *rpoB, pnp*, and *spoT*. Mutations in these three genes have already been associated with antibiotic resistance in other studies (58) or were selected in comparable evolution experiments with *H. influenzae* (43,54), *E. coli* (59), and *Staphylococcus aureus* (60). RNA polymerase subunit β is essential for transcription and even though specific mutations in *rpoB* are known to cause rifampicin resistance, the gene was also recently associated with beta-lactam susceptibility in *Neisseria gonorrhoeae* (27), *S. aureus* (27), and *Bacillus subtilis* (29). The highly conserved PNPase, which influences mRNA and small RNA stability, is described to affect resistance and tolerance to ciprofloxacin and aminoglycosides in *Pseudomonas aeruginosa*, respectively (30,31). The pyrophosphohydrolase encoded by *spoT* synthesizes and hydrolyzes alarmone ppGpp, which induces the expression of stress resistance components, and is involved in resistance and tolerance against different antibiotics including beta⍰lactams for example in *P. aeruginosa* and *Helicobacter pylori* (61– 63). In our study, the effect of mutations in *rpoB, pnp* and *spoT* genes was limited to a two-fold MIC increase. Therefore, one of the detected mutations alone does not result in beta⍰lactam resistance. However, these novel association with decreased beta-lactam susceptibility in *H. influenzae* highlight the complex interplay of various genetic factors in resistance mechanisms.

Drug resistance mutations often entail a reduced fitness in the absence of antibiotic pressure (64,65) as the antibiotic target is often involved in important cellular processes (66). Indeed, almost all of the clones examined exhibited a significant fitness deficit, with significant variability across mutants. Notably, mutations associated with a higher increase in MIC do not necessarily entail a greater fitness deficit. This is exemplified by the amino acid substitution p.Asp551Tyr in PBP3, which resulted in only a moderate fitness decrease of approximately 9% (within the wild-type genomic background) but was associated with an 8-fold increase of the MIC of ceftriaxone. Fitness levels can influence the evolutionary success of a mutant. It is assumed that mutants with little or no fitness deficit tend to be more successful (67). This is supported by studies on *Mycobacterium tuberculosis* and *Staphylococcus aureus*, which discovered mutations with the lowest fitness costs to be the ones most frequently identified in resistant clinical isolates (68–70).

In addition, the frameshift mutation in *ompP2* was associated with a relatively moderate fitness deficit of approximately 12%. At the same time, this mutation provided a significant advantage in the presence of the antibiotic, with a 23.5-fold increase in the MIC of ceftriaxone. In the evolution experiment, the *ompP2* frameshift mutations outcompeted the inversion mutant over time, as both mutations were initially detected in the second passage, but just the frameshift mutation persisted until passage 20. However, applied bottlenecks, such as the 1:100 dilution used for passaging in this study, inherently impose a random selection, as only a small, randomly chosen subset of the growing population can proliferate in the next passage, thus increasing the effect of genetic drift. Genetic drift influences the manifestation of specific mutations, as certain variants may be lost or disproportionately represented purely by chance (71–73). A specificity of OmpP2 for NAD as well as highly reduced uptake of NAD in *ompP2* knockout strains has been described for *H. influenzae* (74). This might explain the reduced fitness observed for clones with mutated *ompP2*, as *H. influenzae* is dependent on extracellular NAD (75) due to the lack of enzymes necessary for de novo biosynthesis (76).

In cases of entailed fitness costs, fitness can be restored by reverting the resistance causing mutation, which is just possible in the absence of antibiotic pressure (77), and by the acquisition of compensatory mutations (64). We identified that the p.Thr443Ala substitution in PBP3 mitigates the fitness deficit caused by p.Ala530Ser in PBP3 (and p.Val423fs in *pnp*), while this amino acid substitution is also associated with a slight increase in the MIC of third-generation cephalosporins. Such mutations likely present an increased risk for being selected in the presence and the absence of the antibiotic.

Resistance-conferring mutations can additionally affect susceptibility to other antibiotics. Simultaneously acquired resistance to another antibiotic is referred to as cross-resistance, while collateral sensitivity describes an evolutionary trade-off where antibiotic resistance to one antibiotic confers increased susceptibility to another antibiotic. Robust collateral sensitivity phenotypes could help to develop new evolutionary-based treatment strategies and therefore may reduce resistance emergence (78).

We observed collaterally increased MICs of fluoroquinolones in *ompP2* mutated clones with decreased beta-lactam susceptibility. This influence of OmpP2 on fluoroquinolone susceptibility in *H. influenzae* is supported by evolution experiments with clinical isolates of *H. influenzae* in the presence of moxifloxacin that selected for *ompP2* mutations and MICs exceeding CLSI breakpoints for moxifloxacin (54). In our experiments MICs did not exceed EUCAST clinical breakpoints for moxifloxacin (> 0.125 mg/L) and levofloxacin (> 0.06 mg/L). However, the success of treatment with fluoroquinolones might be reduced in these cases. For the macrolide clarithromycin we observed differential effects in the *ompP2* mutants. These observations may be clinically relevant, as *H. influenzae* could be treated with co-trimoxazole, macrolides, fluoroquinolones, or tetracycline in cases of known resistance or intolerance to β-lactam antibiotics (79).

Overall, our study identified the emergence of collateral sensitivity towards the three tested aminoglycosides, against the macrolide clarithromycin, the glycopeptide vancomycin, and also the polymyxin drug colistin. Especially the reduced MIC to vancomycin in one evolution experiment was surprising as this antibiotic is only active against gram positive bacteria. This effect might be linked to the detected mutation in the *mltC* gene, as increased susceptibility to vancomycin under acidic pH has been observed in a MltC/MltE deletion mutant of *Escherichia coli* (80). Overall, these patterns of evolved collateral sensitivity were observed for more than 50% of the tested clones. Even though they did not emerge in all cases, collateral sensitivities seem to occur frequently towards specific antibiotics and may thus be a promising focus for optimizing antibiotic treatment designs.

The therapeutic potential of such evolutionary trade-offs clearly deserves further study, especially using combinatorial and sequential treatments with a beta-lactam and an aminoglycoside or clarithromycin. Unfortunately, due to the ototoxicity and nephrotoxicity, which are common side effects of aminoglycosides, a potential combinatorial or alternating treatment would not be appropriate for simple respiratory tract infections caused by *H. influenzae* (81), but rather of value for severe infections like meningitis or sepsis. In fact, beta-lactam–aminoglycoside combinations are used in cases like neonatal meningitis or severe sepsis of unknown etiology, where *H. influenzae* is only rarely the causative pathogen. Nevertheless, in such cases, according to our findings this therapy could be effective.

### Limitations

*H. influenzae* Rd KW20, a well-characterized reference strain, was used as the starting point for all evolution experiments. This represents a limitation, as evolutionary trajectories and phenotypic effects of mutations may vary by genetic background (33). While our findings offer a valuable foundation, the generalizability of resistance-associated fitness and collateral effects should be confirmed in diverse clinical isolates. Future studies incorporating the detection of resistance mutations identified here will be essential to assess their broader clinical relevance across diverse genetic backgrounds.

## Conclusion

Overall, this study revealed the fast evolution of beta-lactam resistance in the opportunistic human pathogen *H. influenzae*. Evolved resistance is mediated by an accumulation of mutations in known target genes and in two populations associated with the emergence of heterogeneity in susceptibility. Almost all resistant clones suffered from reduced competitive fitness in the absence of antibiotics and often expressed collateral sensitivity towards other drug classes, especially aminoglycosides. These findings indicate robust evolutionary trade-offs that are caused by the resistance-conferring mutations and may represent a promising focus for optimizing antibiotic treatment designs. Similarly, a further analysis of novel resistance mutations and of epistatic effects may additionally assist to improve the development of new antimicrobial treatment strategies.

## Acknowledgments

We thank T. Niemann, V. Mohr, M. Mundzeck, T. Struve from the Research Center Borstel, Germany for excellent technical assistance.

## Transparency Declaration

The authors declare that they do not have any competing interests.

## Author contributions

SP, MD and MM conceived the study. SP performed evolutionary experiments, phenotyping and statistical analysis. MM performed Bayesian clock dating analysis. SP, MD, and CU performed next generation sequencing. SP and MM wrote the first draft, all authors reviewed the manuscript, and provided intellectual input. All authors read and approved the final manuscript.

## Supplementary Figures

Supplementary Figure S1: Graphical protocol of the performed multi-step evolution experiment (generated with BioRender).

Supplementary Figure S2: Graphical protocol of the performed competitive growth assay (generated with BioRender).

Supplementary Figure S3: Maximum likelihood phylogeny of *H. influenzae* Rd KW20 clones evolved in the absence of an antibiotic pressure (A, 40 clones) and in three replicate populations evolved in ceftriaxone (B, replicate 1 with 57 clones; C replicate 2 with 56 clones; D, replicate 3 with 59 clones). Amino acid substitutions in the penicillin binding protein 3 (PBP3) and presence of mutations in specified genes as well as minimum inhibitory concentration (MIC) changes over time are color coded. Strains that exceeded the clinical breakpoint for ampicillin/cefotaxime/ceftriaxone according to EUCAST breakpoints are marked with a white star at the MIC heatmap.

The specified genes are those in which at least 20 clones from the whole set of selected clones exhibited mutations but did not mutate under antibiotic free conditions.

Supplementary Figure S4: Crystal structure of Outer membrane protein P2 of wild-type *H. influenzae* Rd KW20 (A) and mutated OmpP2 (B) truncated due to p.Thr126fs (c.376_377insT). Structures were predicted using the AlphaFold Server powerd by AlphaFold 3 (26) and visualized using PyMOL v3.0. Deviations between the two variants are highlighted in red.

Supplementary Figure S5: Crystal structure of Outer membrane protein P2 of wild-type *H. influenzae* Rd KW20, shown in side view (A) and bottom view (B) and two variants of OmpP2 altered due to p.Gly354fs (C and D, inversion) and p.Val353fs (E and F, deletion). Structures were predicted using the AlphaFold Server powerd by AlphaFold 3 (26) and visualized using PyMOL v3.0. Deviations between the two variants are labeled and highlighted in yellow (hydrophobic), green (polar uncharged), and blue (positively charged).

Supplementary Figure S6: Gradient diffusion test demonstrating phenotypic heterogeneity. The image shows two distinct inhibition zones surrounding the antibiotic gradient strip, indicative of a heterogeneous bacterial population. The inner zone corresponds to a subpopulation with higher antibiotic resistance, while the outer zone represents the more susceptible majority.

