## Supplementary figures and images for "Evolution of beta-lactam resistance causes fitness reductions and several cases of collateral sensitivities in the human pathogen *Haemophilus influenzae*"

### Supplementary Figure S1

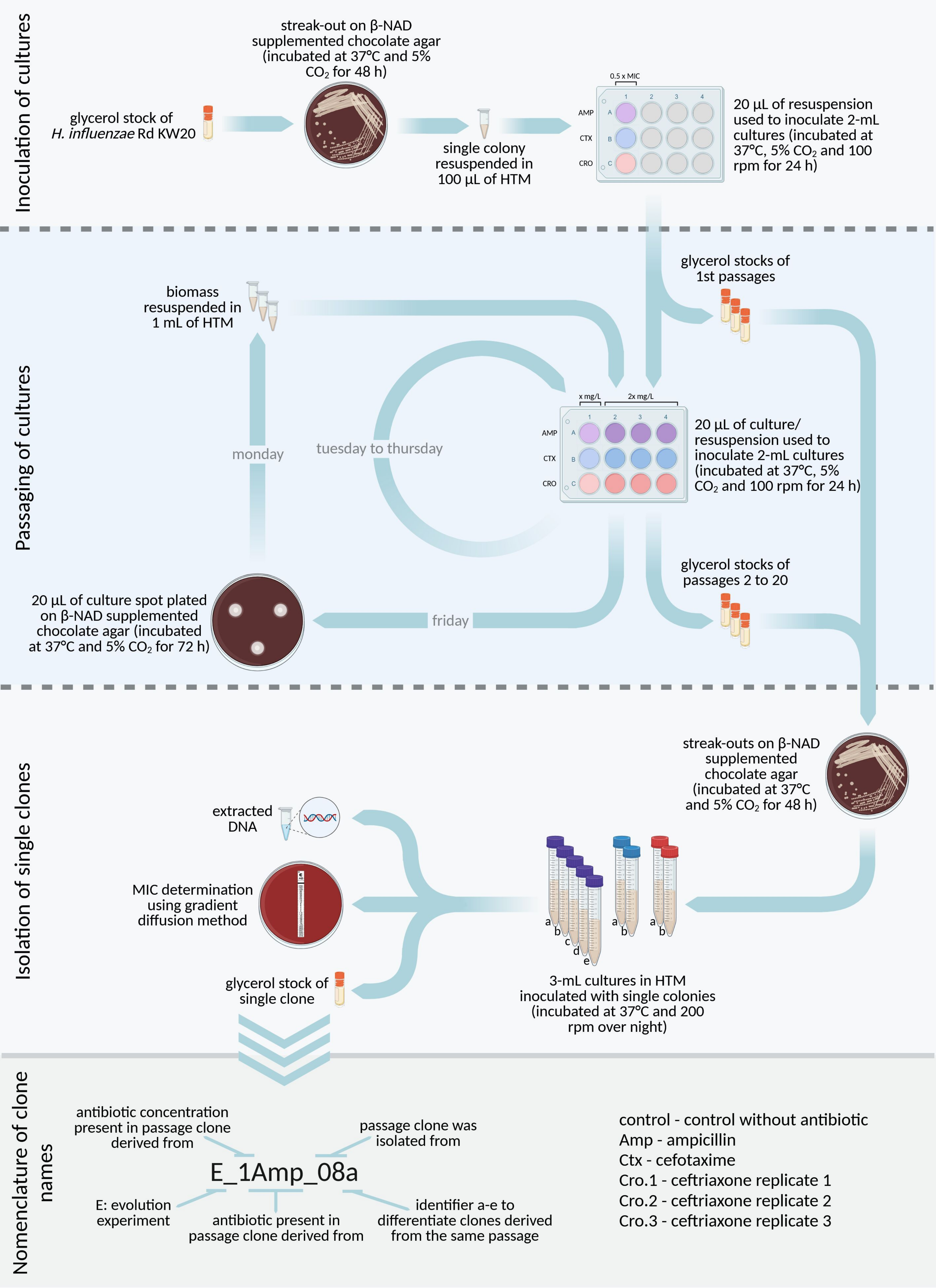

### Supplementary Figure S2

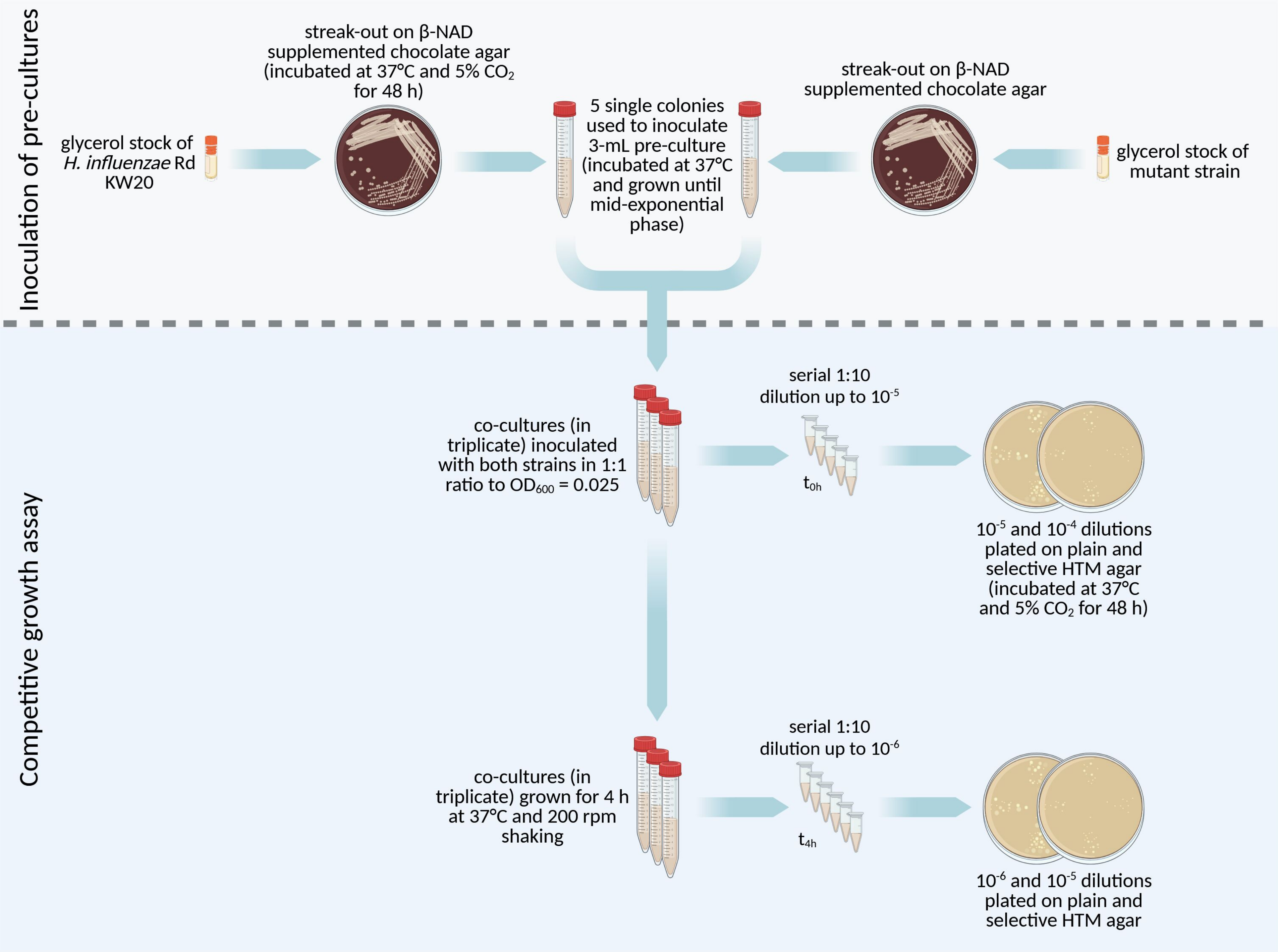

### Supplementary Figure S3

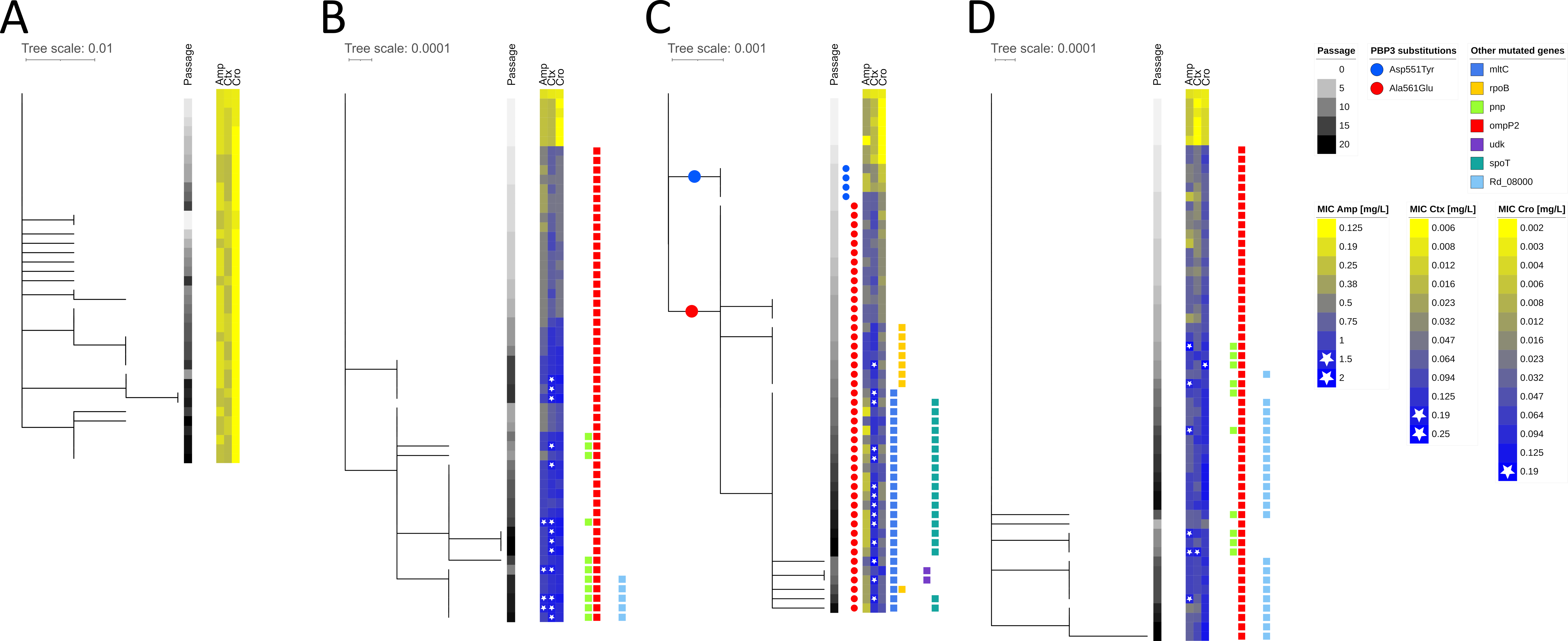

### Supplementary Figure S4

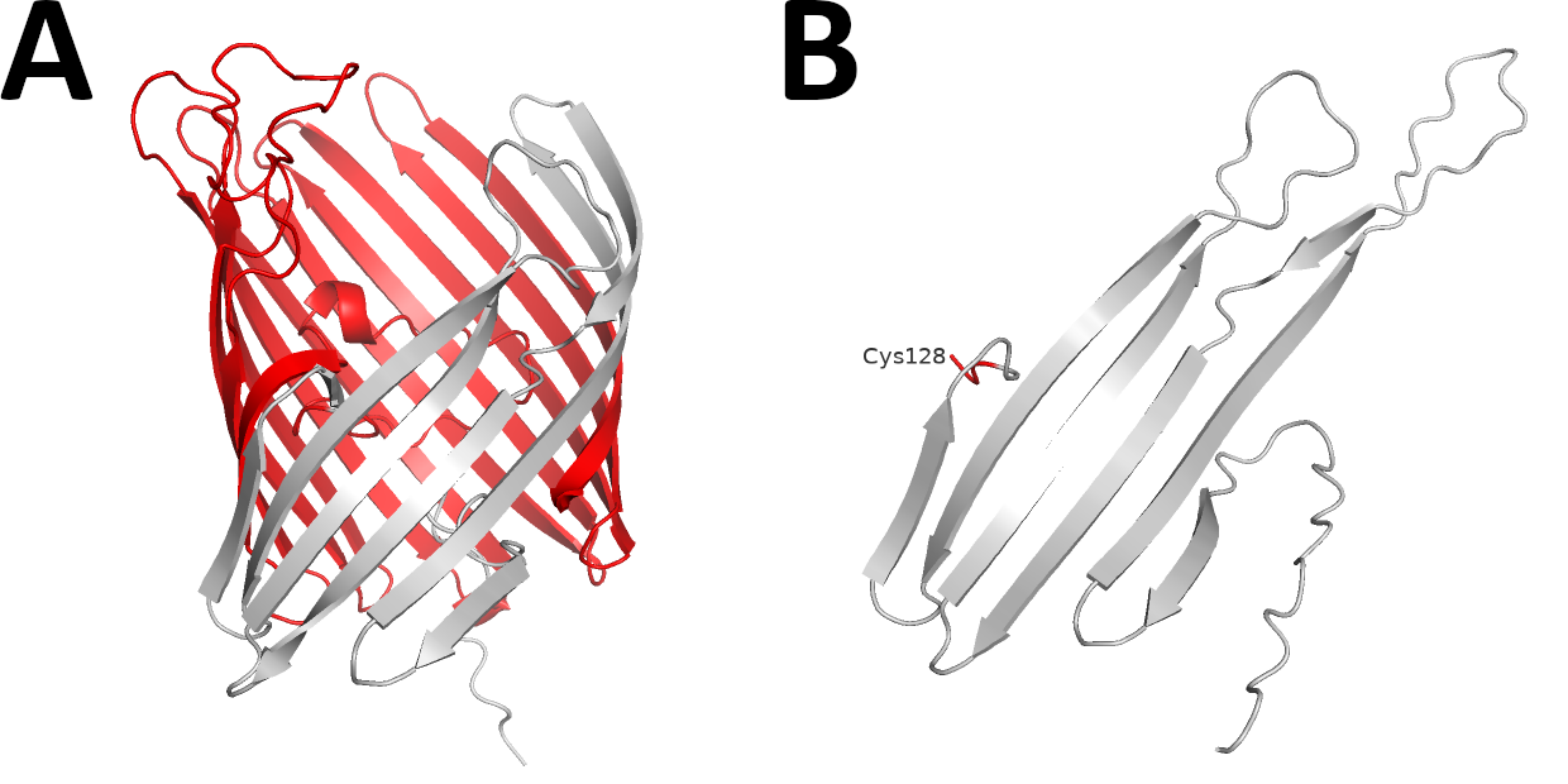

### Supplementary Figure S5

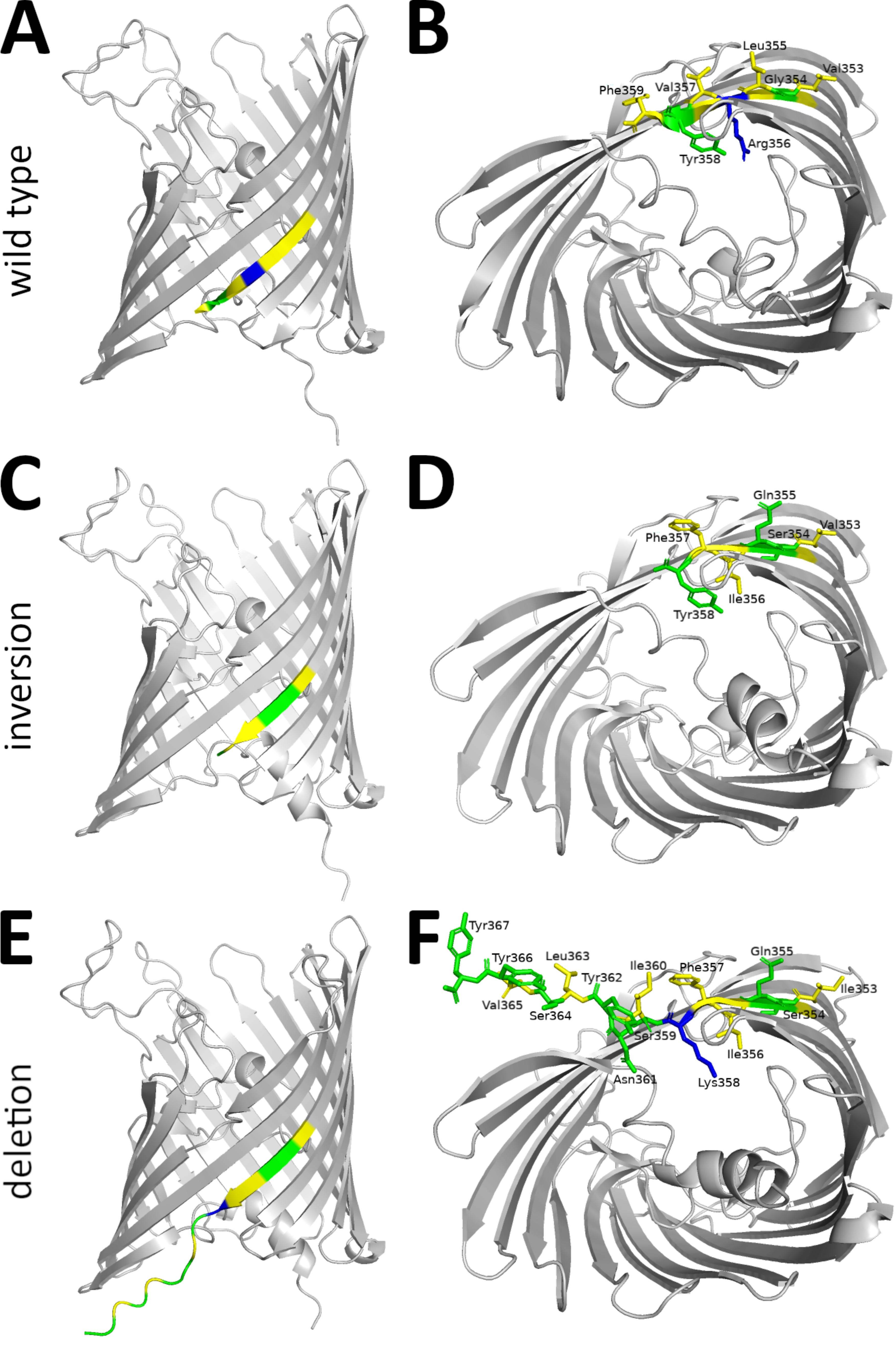

### Supplementary Figure S6

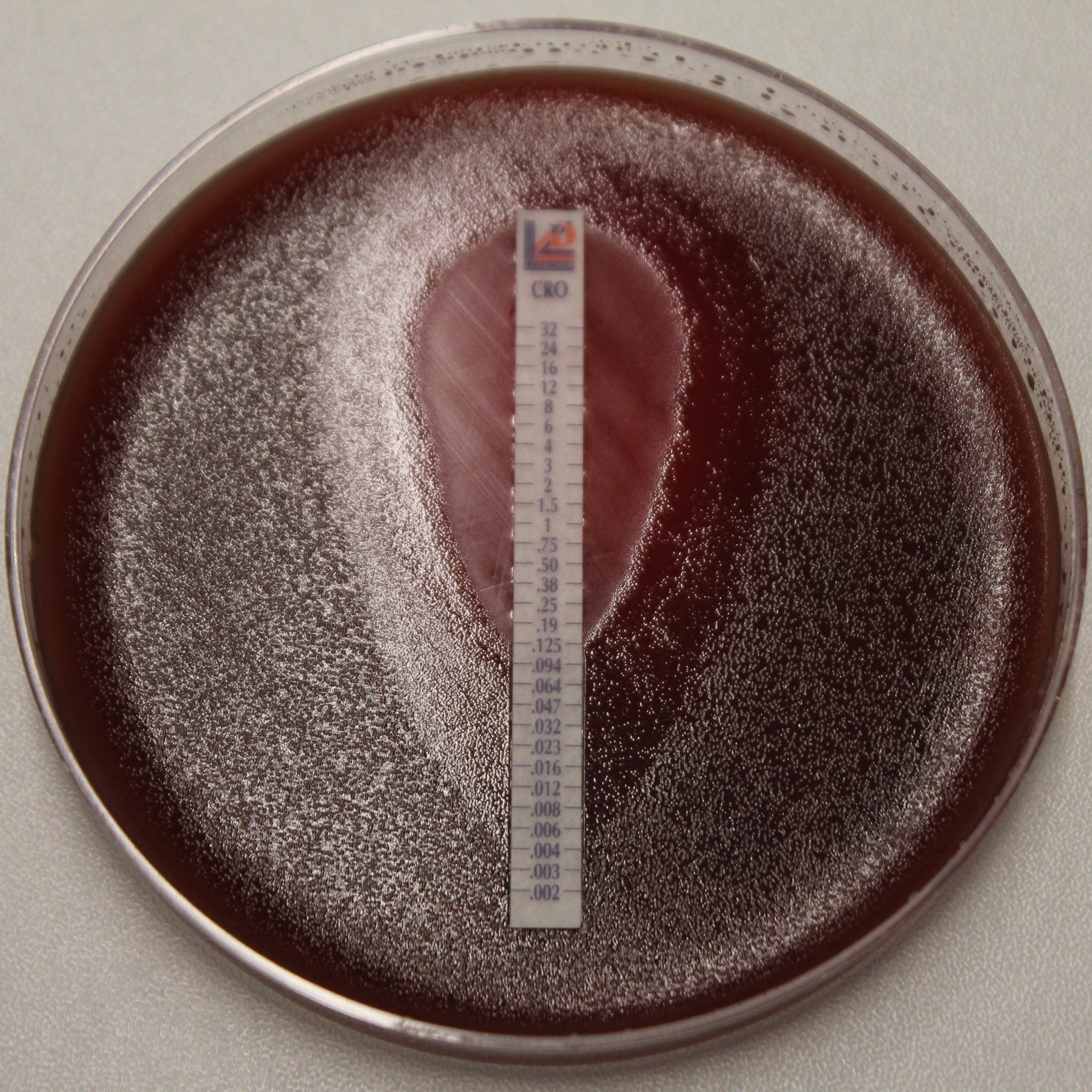
